# Choice seeking is motivated by the intrinsic need for personal control

**DOI:** 10.1101/2022.09.20.508669

**Authors:** Jérôme Munuera, Marta Ribes Agost, David Bendetowicz, Adrien Kerebel, Valérian Chambon, Brian Lau

## Abstract

When deciding between options that do or do not lead to future choices, humans often choose to choose. We studied choice seeking by asking subjects to decide between a choice opportunity or performing a computer-selected action. Subjects preferred choice when these options were equally rewarded, even deterministically, and were willing to trade extrinsic rewards for the opportunity to choose. We explained individual variability in choice seeking using reinforcement learning models incorporating risk sensitivity and overvaluation of rewards obtained through choice. Degrading perceived controllability diminished choice preference, although willingness to repeat selection of choice opportunities remained unchanged. In choices following these repeats, subjects were sensitive to rewards following freely chosen actions, but ignored environmental information in a manner consistent with a desire to maintain personal control. Choice seeking appears to reflect the intrinsic need for personal control, which competes with extrinsic reward properties and external information to motivate behavior.

**Author summary:** Human decisions can often be explained by the balancing of potential rewards and punishments. However, some research suggests that humans also prefer opportunities to choose, even when these have no impact on future rewards or punishments. Thus, opportunities to choose may be intrinsically motivating, although this has never been experimentally tested against alternative explanations such as cognitive dissonance or exploration. We conducted behavioral experiments and used computational modelling to provide compelling evidence that choice opportunities are indeed intrinsically rewarding. Moreover, we found that human choice preference varied according to individual risk attitudes, and expressed a need for personal control that competes with maximizing reward intake.

---

Preference for choice has been observed in humans(1–6) as well as other animals including rats(7), pigeons(8) and monkeys(9,10). This free-choice premium can be behaviorally measured by having subjects perform trials in two stages: a decision is first made between the opportunity to choose from two terminal actions (*free*) or to perform a mandatory terminal action (*forced*) in the second stage(7). Although food or fluid rewards follow terminal actions in non-human studies, choice preference in humans can be elicited using hypothetical outcomes that are never obtained(3,11). Thus, choice opportunities appear to possess or acquire value in and of themselves. It may be that choice has value because it represents an opportunity to exercise control, which is itself intrinsically rewarding(1,4,12). Personal control is central in numerous psychological theories, where constructs such as autonomy(13,14), controllability(15,16), personal causation(17), effectance(18), perceived behavioral control(19) or self-efficacy(15) are key for motivating behaviors that are not economically rational or easily explained as satisfying basic drives such as hunger, thirst, sex, or pain avoidance(20).

There are alternative explanations for choice seeking. For example, subjects may prefer choice because they are curious and seek information(21,22), or they wish to explore potential outcomes to eventually exploit their options(23), or because they seek variety to perhaps reduce boredom(24) or keep their options open(3). By these accounts, however, the expression of personal control is not itself the ends, but rather a means for achieving an objective that once satisfied reduces choice preference. For example, choice preference should decline when there is no further information to discover in the environment, or after uncertainty about reward contingencies have been satisfactorily resolved.

Choice seeking may also arise due to selection itself altering outcome representations. Contexts signaling choice opportunities may acquire distorted value through choice-induced preference change(25). By this account, deciding between equally valued terminal actions generates cognitive dissonance that is resolved by post-choice revaluation favoring the chosen action(25,26). This renders the free option more valuable than the forced option since revaluation only occurs for self-determined actions(27,28). Alternatively, subjects may develop distorted outcome representations through a process related to the winner’s or optimizer’s curse(29), whereby optimization-based selection upwardly biases value estimates for the chosen action. One algorithm subject to this bias is Q-learning(30), where action values are updated using the maximum value to approximate the maximum expected value. In a two-stage task, the free action value is biased upwards due to considering only the best of two possible future actions, while the forced action value remains unbiased since there is only one possible outcome(31). Again, the expression of personal control is not itself the ends for these selection-based accounts, and both predict that choice preference should be reduced when terminal rewards associated with the free option are clearly different.

Data from prior studies does not arbitrate between competing explanations for choice-seeking. Here, we used behavioral manipulations and computational modelling to explore the factors governing human preference for choice. In the first experiment, we altered the reward contingencies associated with terminal actions in order to rule out curiosity, exploration, variety-seeking, and selection-based explanations for choice seeking. In the second experiment, we used a titration procedure to measure the value of choice relative to an extrinsic reward available in the environment (i.e., money). We then used reinforcement learning models to show that optimistic learning (considering the best possible future outcome) was insufficient to explain individual variability in choice seeking. Rather, subjects adopted different *decision attitudes*, the desire to make or avoid decisions independent of the outcomes(11), which were balanced against differing levels of risk sensitivity. Finally, in the third experiment, we sought to test whether choice preference was motivated by personal control beliefs. We manipulated the perceived controllability of the task and found that subjects’ willingness to repeat a free choice was not affected by the lack of objective controllability over reward outcome. Importantly, subjects were sensitive to past rewards only in trials where state outcomes could be attributed to self-determined choice, and ignored rewards on trials where there was an apparent loss of control. Together, our results support the hypothesis that human preference for choice opportunities derives from the intrinsic motivation for personal control.

## Results

Subjects performed repeated trials with a two-stage structure (Fig. 1). In each trial, subjects made a 1^st^-stage choice between two options defining the 2^nd^-stage: the opportunity to choose between two fractal targets (*free*) or performing an obligatory selection of another fractal target (*forced*). Extrinsic rewards (€) were delivered only for terminal (i.e., 2^nd^-stage) actions. If subjects chose the *forced* option, the computer always selected the same fractal target for the subjects. If subjects chose the *free* option, they had to choose between two fractal targets associated with two different terminal states. We fixed reward contingencies in blocks of trials, and used unique fractal targets for each block. We divided each block into an initial training phase (Fig. 1B) followed by a test phase (Fig. 1C) to ensure that the subjects learned the associations between the different fractal targets and extrinsic reward probabilities.

**Figure 1.**
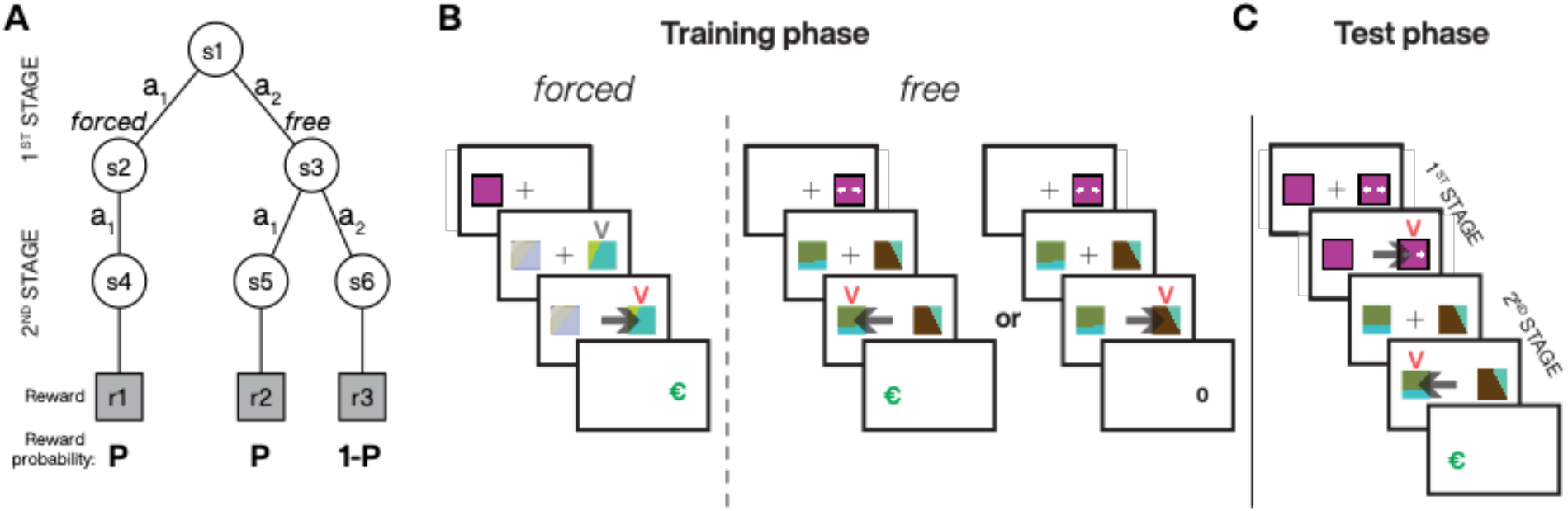
Two-stage task structure. **A**. State diagram illustrating the 6 possible states (s), actions (a) and associated extrinsic reward probabilities (e.g., P = 0.5, 0.75 or 1 for blocks 1 to 3, respectively); s2 and s3 were represented by two different 1^st^-stage targets (e.g., colored squares with or without arrows for *free* and *forced* trials, respectively) and s4 to s6 were associated to three different 2^nd^-stage targets (fractals). **B**. Sequence of events during the training phase where the subjects learned the contingencies between the fractal targets and their reward probabilities (P) associated with the *forced* (no choice) and *free* (choice available) options. When training the reward contingencies associated with the *forced* option, subjects’ actions in the 2^nd^-stage had to match the target indicated by a grey V-shape and was always the same (s4). When training the reward contingencies associated with the *free* option, no mandatory target is present at the 2^nd^-stage (s5 or s6 can be chosen) but one of the targets is more rewarded when P > 0.5. Black arrows represent the selection of the target by the subject. **C**. Sequence of events during the test phase: subjects first decided between the *free* or *forced* option and then experienced the associated 2^nd^-stage. Rewards, when delivered, were represented by a large green euro symbol (€).

### Free choice preference across different extrinsic reward probabilities

In experiment 1, we varied the overall expected value by varying the probability of extrinsic reward delivery (P) across different blocks of trials. These probabilities ranged from 0.5 to 1 across the blocks (i.e., low to high), and the programmed probabilities in *free* and *forced* 2^nd^-stage rewards were equal (Fig. 2A). For example, in high probability blocks, we set the probabilities of the *forced* terminal action and of one of the *free* terminal actions (a1) to 1, and set the probability of the second *free* terminal action (a2) to 0. Therefore, the maximum expected value was equal for the *free* and *forced* options.

**Figure 2.**
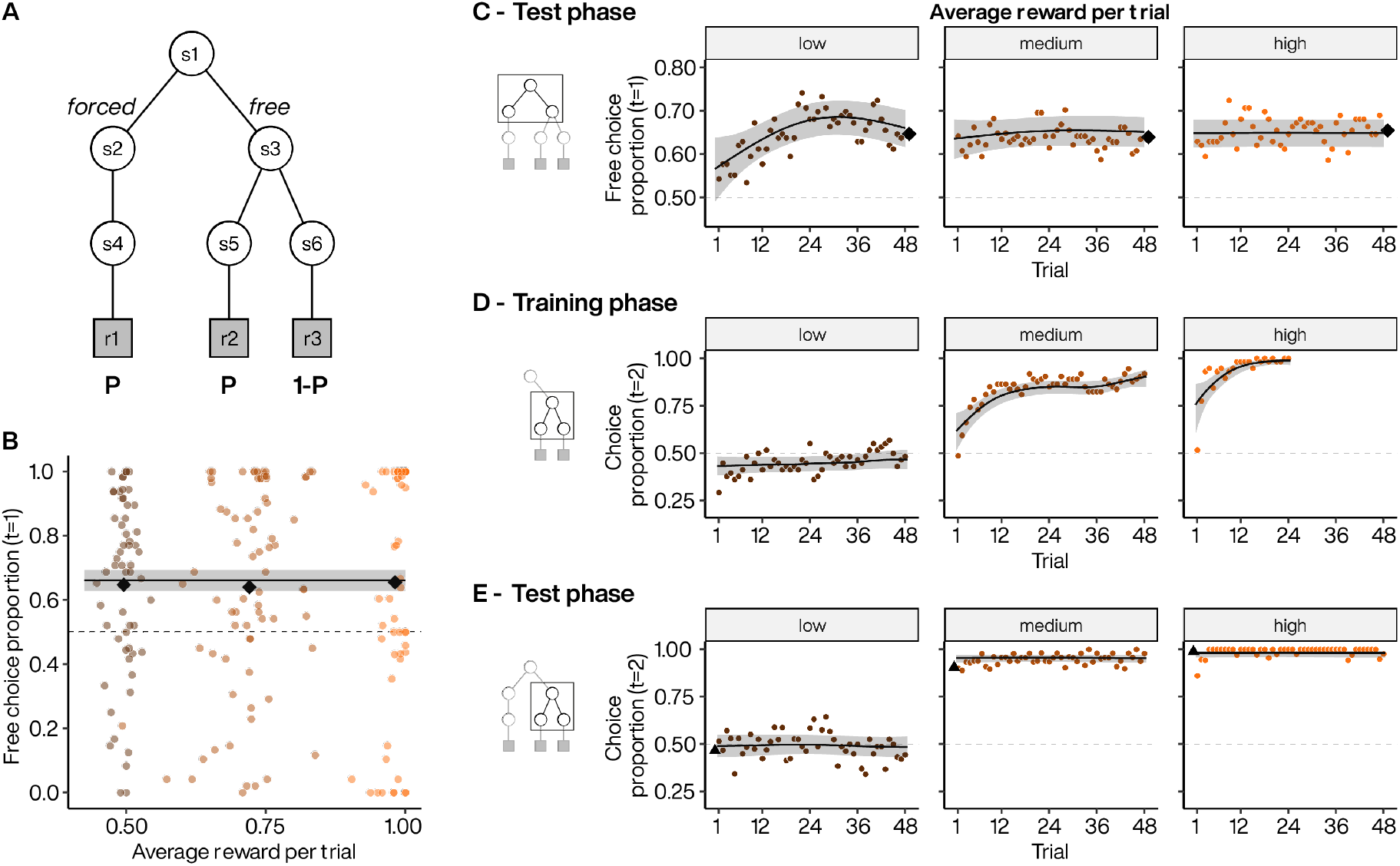
Choice preference across different absolute extrinsic reward probabilities. **A**. Experiment 1 task design where maximal extrinsic reward probabilities increased equally across *free* and *forced* options. **B**. Subject preference for *free* option during 1^st^-stage. Colored points indicate individual subject mean choice preference per block, plotted against the obtained average reward. Black diamonds indicate the average of subject means per block. Line indicates the estimated choice preference from a GLMM, with 95% CI. **C**. Dynamics of *free* option preference across test phase blocks for low (left), medium (middle) and high (right) absolute extrinsic reward probabilities. Each point represents the average *free* option preference as a function of trial within a block. Diamonds: as in B. Lines indicate the estimated choice preference from a GAMM, with 95% CI. **D** to **E**. Dynamics of the selection of the most rewarded 2^nd^-stage targets in *free* option for low (left), medium (middle) and high (right) during the training (D) and test (E) phases. Note that in left panels, the probability of extrinsic rewards is equal for two 2^nd^-stage targets (P=0.5). We labelled the best choice as 1 when P > 0.5. Triangles represents the final average selection at the end of the training phases. Lines: as in C.

Subjects chose to choose more frequently, selecting the *free* option in 64% (n=58) of test trials on average (Fig. 2B). The level of preference did not differ significantly across blocks (*p* = 0.857, low = 65%, medium = 64%, high = 66%). We found that subjects immediately expressed above chance preference for the *free* option (Fig. 2C) despite never having actualized 1^st^-stage choices during training. Looking within a block, we found that subjects’ preference remained constant across trials in medium and high reward probability blocks (*p* = 0.22 and 0.6823 for nonlinear smooth by trial deviating from a flat line, respectively; Fig. 2C, middle and right panels). In low probability blocks, subjects started with a lower choice preference that gradually increased to match that observed in the medium and high probability blocks (*p* = 0.0014 for nonlinear smooth by trial; Fig. 2C left panel). The lower reward probability may have prevented subjects from developing accurate reward representations by the end of the training phase, which may have led to additional sampling of the three 2^nd^-stage targets (two in *free* and one in *forced*) in the beginning of the test phase.

### Second-stage performance following *free* selection

We investigated participants’ 2^nd^-stage choices following *free* selection to exclude the possibility that choice preference arose because reward contingencies had not been learned. During the training phase, when P>0.5, participants quickly learned to choose the most rewarded fractal targets (at P=0.5, all fractal targets were equally rewarded) (Fig. 2D). During the test phase, participants continued to select the same targets (Fig. 2E), confirming stable application of learned contingencies (*p* > 0.1 for nonlinear smooth by trial deviating from a flat line for all blocks).

Choice preference was not explained by subjects obtaining more extrinsic rewards following selection of *free* compared to *forced* options. Obtained reward proportions were not significantly different in the low (following selection of *free* vs. *forced*, 0.516 vs. 0.536, *p* = 0.276) or medium (0.746 vs. 0.762, *p* = 0.322) probability blocks. In contrast, in high probability blocks, subjects received significantly fewer rewards on average after *free* selection than after *forced* selection (0.989 vs. 1, *p* = 0.0016). In this block, reward was fully deterministic, and *forced* selection always led to a reward, whereas *free* selections could lead to missed rewards if subjects chose the incorrect target.

### Trading extrinsic rewards for choice opportunities

Since manipulating the overall expected reward did not alter choice seeking behavior at the group-level, we investigated the effect of changing the relative expected reward between 1^st^-stage options. In experiment 2, we tested a new group of 36 subjects for whom we decreased the objective value of the *free* versus *forced* options. This allowed us to assess the point at which these options were equally valued and potentially reversed to favor the initially non-preferred (*forced*) option (Fig. 3A). Thus, we titrated the value of choice opportunity by increasing the reward probabilities following *forced* selection (block 1: P_*forced*_ = 0.75; block 2: P_*forced*_ = 0.85; block 3: P_*forced*_ = 0.95), while keeping the reward probabilities following *free* selection fixed (P_*free*_|a1 = 0.75, P_*free*_|a2 = 0.25 for all blocks).

**Figure 3.**
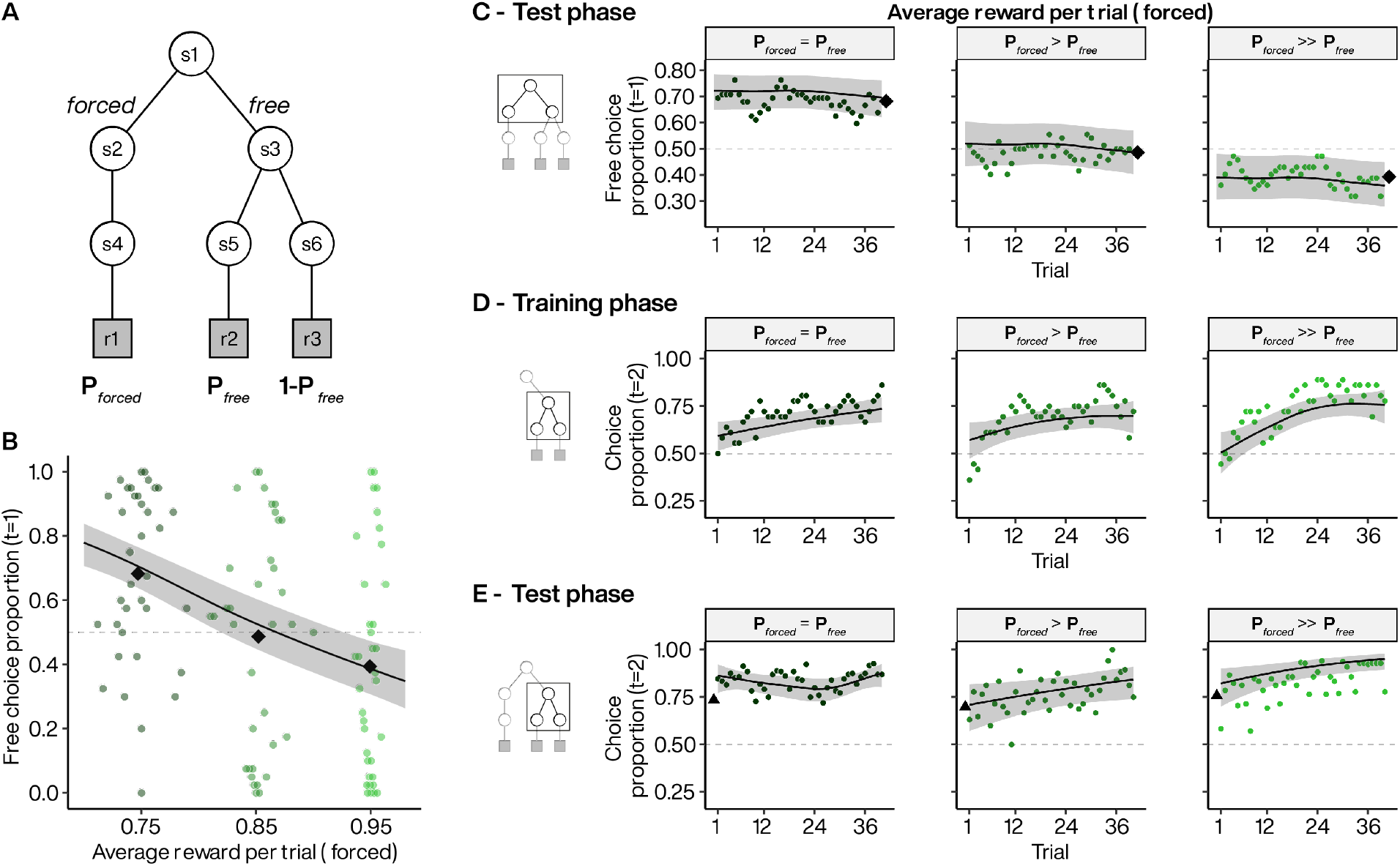
Choice preference across different relative extrinsic reward probabilities. **A**. Experiment 2 task design where extrinsic reward probably is always at P = 0.75 for the highly rewarded target in *free* options but vary from 0.75 to 0.95 across 3 blocks for *forced* options. **B**. Subject preference for *free* option during 1^st^-stage. Colored points indicate individual subject mean choice preference per block, plotted against the average reward in *forced* option. Black diamonds indicate the average of subject means per block. Line indicates the estimated choice preference from a GLMM, with 95% CI. **C**. Dynamics of *free* option preferences across test phase blocks when extrinsic reward probabilities of *forced* options were set at 0.75 (left), 0.85 (middle) and 0.95 (right). Each point represents the average *free* option preference as a function of trial within a block. Diamonds: as in B. Lines indicate the estimated choice preference from a GAMM, with 95% CI. **D to E**. Dynamics of the selection of the most rewarded 2^nd^-stage targets in *free* option when extrinsic reward probabilities of *forced* options are set at 0.75 (left), 0.85 (middle) and 0.95 (right) during the training (D) and test (E) phases. Triangles represents the final average selection at the end of the training phases. Lines: as in C.

As in experiment 1, we found that subjects preferred choice when the extrinsic reward probabilities of the *free* and *forced* options were equal (block 1: 68% 1^st^-stage choice in favor of *free*; Fig. 3B, dark green). Increasing the reward probability associated with the *forced* option significantly reduced choice preference (*p* = 0.00344, Fig. 3B) to 49% (block 2) and 39% (block 3). We estimated the population preference reversal point at P_*forced*_ = 0.88, indicating that indifference was obtained on average when the value of the *forced* option was 17% greater than that of the *free*. We found that subjects’ preference remained constant across trials when reward probabilities were equal (*p* = 0.875 for nonlinear smooth by trial; Fig. 3C, left panel). Although reduced overall, the selection of the *free* option also did not vary across trials in blocks 2 and 3 (*p* = 0.737 and 0.078 for nonlinear smooth by trial, respectively). Furthermore, as in experiment 1, subjects acquired preference for the most rewarded 2^nd^-stage targets during the learning phase (Fig.3D) and continued to express this preference during the test phase in all three blocks (Fig. 3E). Thus, the decrease in choice preference was not related to failure to learn the reward contingencies during the training phase.

Although decreasing the relative value of the *free* option reduced choice preference, most subjects did not switch exclusively to the *forced* option. Even in block 3, where the *forced* option was set to be rewarded most frequently (P_*forced*_ = 0.95 versus P_*free*_ = 0.75), 32/36 subjects selected the *free* option in a non-zero proportion of trials. Since exclusive selection of the *forced* option would maximize extrinsic reward intake, continued *free* selection indicates a persistent appetency for choice opportunities despite their diminished relative extrinsic value.

### Reinforcement-learning model of choice seeking

We next sought to explain individual variability in choice behavior using a value-based decision-making framework. We first used mixed logistic regression to examine whether rewards obtained from 2^nd^-stage actions influenced 1^st^-stage choices. We found that obtaining a reward on the previous trial significantly increased the odds that subjects repeated the 1^st^-stage selection that ultimately led to that reward (*p* < 0.0001, odds ratio rewarded/unrewarded on previous trial: 1.92 ±95% CI [1.40, 2.60]). This suggest that subjects continued to update their extrinsic reward expectation based on experience during the test phase. We therefore leveraged the framework of temporal-difference reinforcement learning (TDRL) to provide a model-based characterization of the emergence of choice preference.

We fitted TDRL models to individual data using two distinct features to capture individual variability across different extrinsic reward contingencies. The first feature was a free choice bonus added to self-determined actions as an intrinsic reward. This can lead to overvaluation of the *free* option via standard TD learning. The second feature modifies the form of the future value estimate used in the TD value iteration, which in common TDRL variants is, or approximates, the best future action value (Q-learning or SARSA with softmax behavioral policy, respectively). We treated both Q-learning and SARSA together as optimistic algorithms since they are not highly discriminable with our data (Supplementary Fig. 1). We compared this optimism with another TDRL variant that explicitly weights the best and worst future action values (Gaskett’s *β*-pessimistic model(32)), which could capture avoidance of choice opportunities through increased weighting of the worst possible future outcome (pessimistic risk attitude). For example, risk is maximal in the high reward probability block in experiment 1 since selection of one 2^nd^-stage target led to a guaranteed reward (best possible outcome) whereas selection of the other target led to guaranteed non-reward (worst possible outcome).

We found that it was necessary to incorporate the overvaluation of rewards obtained from *free* actions to predict choice preference in experiment 1 (Fig. 4A). Moreover, the magnitude of the bonus was significantly associated with increasing choice preference during the 1^st^-stage of the trials (*p* = 0.0005 for nonlinear smooth, Fig. 4B). Therefore, optimistic or pessimistic targets alone were insufficient to explain individual choice preference across different extrinsic reward contingencies. We found that a pessimistic target best fitted about 28% (16 of 58) of the subjects in experiment 1. Moreover, most pessimistic subjects (13 of 16) were best fitted with a model including a free choice bonus to balance risk and decision attitudes across reward contingencies. In experiment 1, we introduced risk by varying the difference in extrinsic reward probability for the best and worst outcome following *free* selection. The majority of so-called ‘pessimistic subjects’ preferred choice when extrinsic reward probabilities were low, but their weighting of the worst possible outcome decreased this preference as risk increased (Fig. 4C, pink). Thus, pessimistic subjects avoided the *free* option despite rarely or never selecting the more poorly rewarded 2^nd^-stage target during the test phase.

**Figure 4.**
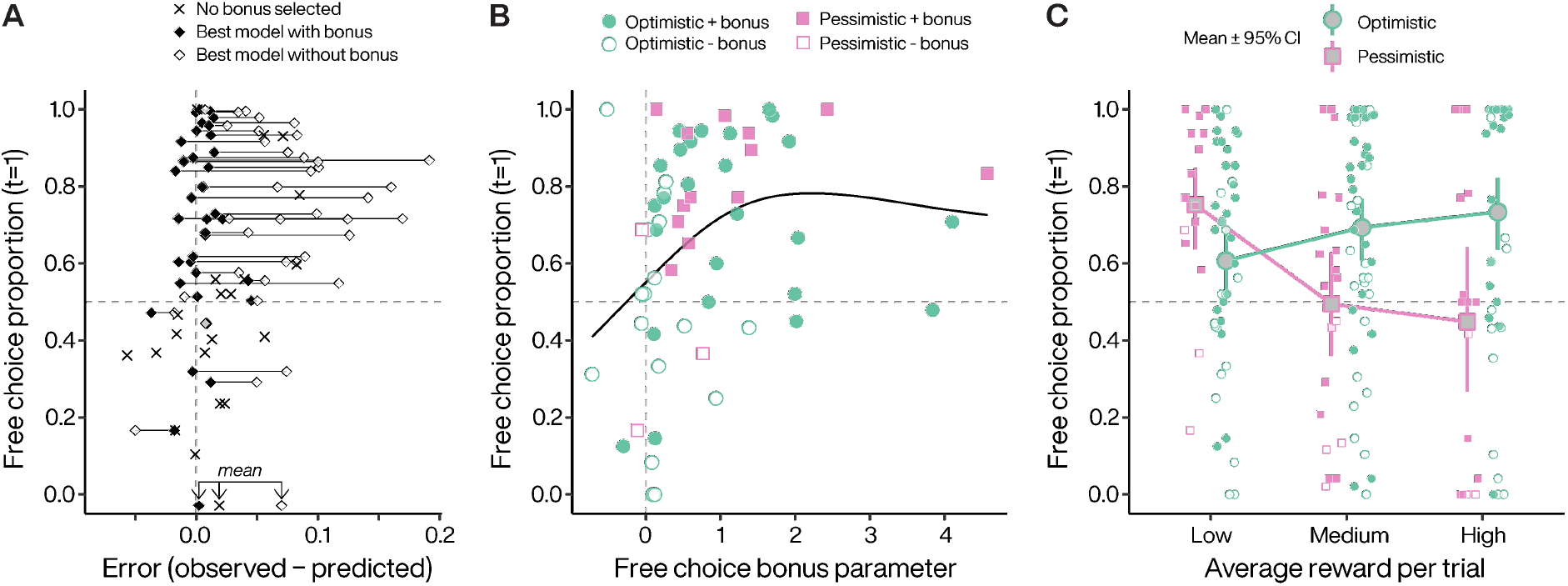
Reinforcement learning models capture individual choice behavior. **A**. Obtained free choice proportion as a function of model error in experiment 1, averaged over all conditions. For subjects where the selected model did not include a free choice bonus, only one symbol (X) is plotted. For subjects where the selected model included a free choice bonus, two symbols are plotted. Filled symbol represents the fit error with the selected model, and the open symbol represents the next best model that did not include a free choice bonus. Lines connect individual subjects. **B**. Bonus coefficients increase as a function of subjects’ preference for *free* options irrespectively of the target policy they used when performing the task. Choice preference from low probability blocks (P=0.5). Filled circles indicate that the best model included a free choice bonus parameter. Line illustrates a generalized additive model smooth. **C**. Pessimistic subjects significantly decrease their *free* option preference as a function of extrinsic reward probabilities. Symbol legend from B applies to the small points representing individual means in C. Error bars for 95% CI.

We also fitted the TDRL variants to individual data from experiment 2, and found that a free choice bonus was also necessary to explain choice preference across extrinsic reward contingencies in that experiment. Four subjects (of 36) were best fitted using the *β*-pessimistic target (see Supplementary Fig. 2) although this may be a conservative estimate since we did not vary risk in experiment 2.

### Influence of action-outcome coherence on choice seeking behavior

We next asked whether choice preference was related to personal control beliefs. To do so, we manipulated the coherence between an action and its consequence over the environment. In experiment 3, we tested the relationship between preference for choice opportunity and the physical coherence of the terminal action by directly manipulating the perceived controllability of 2^nd^-stage actions. We modified the two-stage task to introduce a mismatch between the subject’s selection of the 2^nd^-stage target and the target ultimately displayed on the screen by the computer (Fig. 5A). We did this by manipulating the probability that a 2^nd^-stage target selected by a subject would be swapped for the 2^nd^-stage target that had not been selected. That is, on coherent trials, a subject selecting the fractal on the right side of the screen would receive visual feedback indicating that the right target had been selected. On incoherent trials, a subject selecting the fractal on the right side would receive feedback that the opposite fractal target had been selected (i.e., the left target). To ensure that all other factors were equalized between the two 1^st^-stage choices, we implemented target swaps following both *free* and *forced* selections by adding an additional state to our task (Fig. 5A). In one block of trials, the incoherence was set to 0 and every subject action in the 2^nd^-stage led to a coherent selection of the second target. In the other blocks, we set incoherence to 0.15 or 0.3, resulting in lower perceived controllability between target choice and target selection (e.g., 85% of the time, pressing the left key will select the left target, and in 15% the right target). We set all of the extrinsic reward probabilities associated with the different fractal targets to P = 0.75. Since all 2^nd^ -stage actions had the same expected value, the experiment was objectively uncontrollable because the probability of reward was independent of all actions(16). Moreover, equal reward probabilities ensured that outcome diversity(33,34), outcome entropy(35), and instrumental divergence(36) did not contribute to choice preference since these were all equal between the *forced* and *free* options.

**Figure 5.**
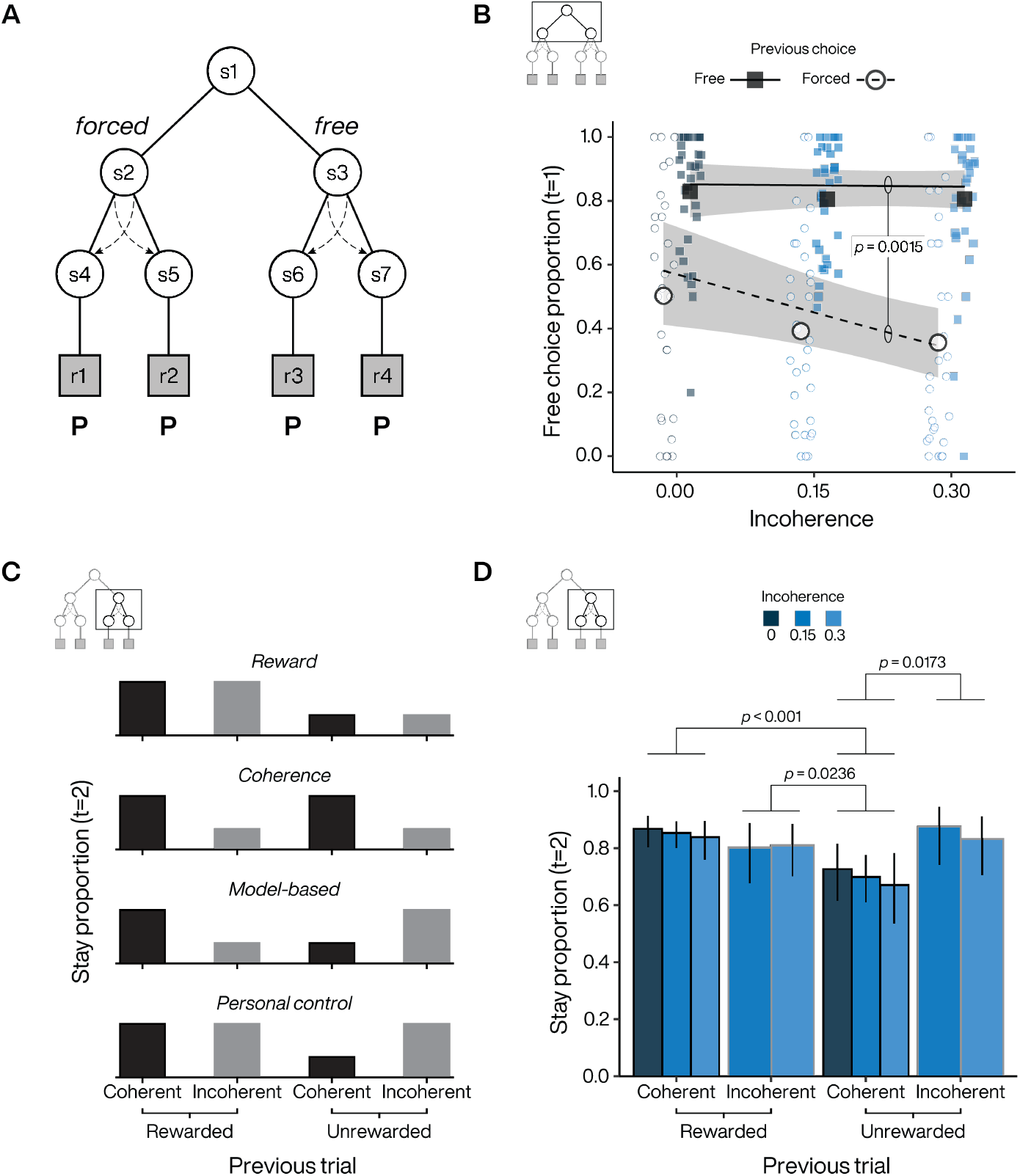
Perceived controllability alters choice preference. **A**. Task design where a 7^th^ state, associated to the *forced* options, has been added to manipulate the incoherence in both *free* and *forced* options. At incoherence = 0, the visual feedback presented to the subject matches their selected target. Extrinsic reward probabilities set at P=0.75 for all the 2^nd^-stage targets. **B**. First-stage probabilities to stay or switch in free options after a *free* and *forced* trial respectively, as a function of the different incoherence blocks. **C**. Second-stage stay probabilities for the different action-state-reward trial type. Each sub-panels represent a putative strategy followed by the subject. **D**. Estimated 2^nd^-stage stay probabilities. Error bars for 95% CI. P-values are displayed for significant pairwise comparisons and adjusted for multiple comparisons.

The same group of participants who performed experiment 2 also performed experiment 3 (n=36). Choice preference was high (70%) in block 1 when coherence was not altered, similar to block 1 from experiment 2 where extrinsic reward was equal between *free* and *forced* options. The only difference between these two blocks was that choosing the *forced* option resulted in the obligatory selection of the same fractal (experiment 2) or one of two fractals randomly selected by the computer (experiment 3), which indicates that subjects’ choice preference was not related to action variability per se following *forced* selection. Moreover, we found that choice preference was significantly correlated (*r* = 0.358, *p* = 0.03175) between block 1 of experiments 2 and 3, highlighting a within-subject consistency in choice preference.

Increasing the incoherence of the 2^nd^-stage actions progressively reduced choice preference (block 2 and 3: 67% and 64% in favor of *free* respectively). As in experiments 1 and 2, choice preference was expressed immediately after the training phase and remained constant throughout the different blocks (Supplementary Fig. 3). We found that the decline in choice preference depended on the 1^st^-stage choice on the previous trial. Specifically, following coherent trials, we found that there was a significant interaction between the previous 1^st^-stage choice (*free* or *forced*) and the degree of incoherence (*p =* 0.0015, Fig. 5B). The difference in slopes was due to decreasing propensity to choose the *free* option following *forced* selection on the previous trial (*p* = 0.0111), with no change in the propensity to choose the *free* option following *free* selection on the previous trial (*p* = 0.8706). Thus, as incoherence increased, subjects tended to stay more with the *forced* option, while maintaining a preference to repeat *free* selections.

The sustained repetition of *free* selections across the different levels of incoherence suggests that subjects may have been seeking to regain control of the environment through self-determined 2^nd^-stage choices. Although the task was objectively uncontrollable since all terminal action-target sequences were associated with the same reward probability, subjects may have developed structure beliefs based on local reward history and target swaps, which could be reflected in 2^nd^-stage patterns of choice. Thus, subjects may have followed a strategy based on reward feedback by repeating only actions associated with a previous reward (illusory maximization of reward intake; Fig.5C, first panel). Alternatively, they could have followed a strategy based on action-outcome incoherence feedback and thus avoided trials associated with a previous target swap (illusory minimization of incoherent states; Fig. 5C, second panel). However, subjects may have also employed another classic strategy known as “model-based” where agents use their (here illusory) understanding of the task structure built from all the information provided by the environment (Fig.5C, third panel)(37). Under this strategy, subjects try to integrate both the reward and target-swap feedback to select the next target in order to maximize reward. For example, an incoherent but rewarded trial would lead to a behavioral switch because the subject has integrated the information provided by the environment (i.e., the target swap induced by the computer), signaling that the other target is actually rewarded (see second bar on third panel of Fig. 5C). Finally, an alternative strategy could rely on maximizing personal (i.e., internal) control, where the subject is the (illusory) agent of the entire sequence of events (i.e., action-state-reward) and would therefore ignore reward outcomes when they are not associated with the selected action-state (Fig.5C, fourth panel).

Results of the stay behavior during 2^nd^-stage choice following *free* selection suggests that subjects seek personal control when choosing between the different fractal targets (Fig.5D). Indeed, when their action was consistent with the state they were choosing (i.e., the coherent fractal target feedback), they took the reward outcome into account to adjust their behavior on the next trial, either by staying on the same target when the trial was rewarded or by switching to the other one when no reward was delivered. However, subjects were insensitive to the reward outcome during incoherent trials as they maintained the same strategy (staying) during subsequent trials, regardless of whether they were previously rewarded or not. This strategy reflects an attempt to regain personal control over the environment at the expense of the task goal of maximizing reward intake.

## Discussion

Animals prefer situations that offer more choice to those that offer less. Although this behavior can be reliably measured using the two-stage task design popularized by Voss and Homzie(7), their conclusion that choice has intrinsic value is open to debate. To rule out alternative explanations for choice-seeking, we performed three experiments in which we clearly separated learning of reward contingencies from testing of choice preference. Our experiments point to a sustained preference for choice opportunities that express an intrinsic need for personal control. Moreover, this need may compete with potentially valuable information for maximizing outcomes or even extrinsic rewards per se.

In the first and second experiments, we varied the reward probabilities associated with terminal actions following *free* and *forced* selection. Consistent with previous studies, subjects preferred the opportunity to make a choice when expected rewards were equal between terminal actions (P = 0.5). Surprisingly, subjects also preferred choice when we increased the value difference between terminal actions in the *free* option, while keeping the *maximum* expected reward equal in the free and forced options (P > 0.5). This sustained preference for choice is potentially economically suboptimal since making a free choice carries the risk of making an error leading to lowered reward intake. The persistence of this preference for free choice even when reward delivery was deterministic (P = 1), makes it unlikely that this preference was due to an underestimation of forced trials due to poor learning of reward contingencies.

Subjects appeared to have understood the reward contingencies as evidenced by their consistent preference for the highest-rewarded 2^nd^-stage fractal, which was acquired during the training phase and expressed during the test phase. This stable 2^nd^-stage fractal selection, together with the immediate expression and maintenance of 1^st^-stage choice preference, renders unlikely accounts based on curiosity, exploration or variety seeking since varying the probability of rewards did not modulate choice preference about two third of the subjects (i.e., optimistic subjects).

Selection-based accounts also have trouble explaining the pattern of results we observed. The idea that post-choice revaluation specifically inflates expected outcomes after choosing the free option can explain choice-seeking when all terminal reward probabilities are equal. However, post-choice revaluation cannot explain choice preference when the terminal reward probabilities in the *free* option clearly differ from one another, since revaluation appears to occur only after choosing between closely valued options(28,38). That is, there is no cognitive dissonance to resolve when reward contingencies are easy to discriminate, and no preference for choice should be observed when the choice is between a surely (i.e., deterministically) rewarded action and a never rewarded action. The existence of choice preference in the deterministic condition (P = 1) also cannot be explained by an optimistic algorithm such as Q-learning, since the maximum action value is equal to the maximum expected value, and the value of the free option is not biased upwards under repeated sampling(31).

Although standard Q-learning could not capture variability across different terminal reward probabilities, we found that combining two novel modifications to TDRL models was able to do so. The first feature was a free choice bonus—a fixed value added to all extrinsic rewards obtained through free actions—that can lead to overvaluation of the free option via standard TD learning. This bonus implements Beattie and colleagues’ concept of *decision attitude*, the desire to make or avoid decisions independent of the outcomes(11). The second feature modifies the form of the future value estimate in the TD value iteration. Zorowitz and colleagues(31) showed that replacing the future value estimate in Q-learning with a weighted mixture of the best and worst future action values(32) can generate behavior ranging from aversion to preference for choice. The mixing coefficient determines how optimism (maximum of future action values, total risk indifference) is tempered by pessimism (minimum of future action values, total risk aversion). In experiment 1, we found that 28% of subjects were best fitted with a model incorporating pessimism, which captured a downturn in choice preference with increasing relative value difference between the terminal actions in the *free* option. Importantly however, individual variability in the TD future value estimates alone did not explain the pattern of choice preference across target reward probabilities, and a free choice bonus was still necessary for most subjects. Thus, the combination of both a free choice bonus (decision attitude) and pessimism (risk attitude) was key for explaining why some individuals shift from seeking to avoiding choice. This was unexpected because the average choice preference in experiment 1 was not significantly different across reward manipulations, highlighting the importance of examining behavior at the individual level. Here, we examined risk using the difference between the best and worst outcomes as well as relative value using probability (see(39)). It may be the case that variability is also observed in how individuals balance the intrinsic rewards with other extrinsic reward properties that can influence choice preference, such as reward magnitude(39).

Why are choice opportunities highly valued? It may be that choice opportunities have acquired intrinsic value because they are particularly advantageous in the context of the natural environment in which the learning system has evolved. Thus, choice opportunities might be intrinsically rewarding because they promote the search for states that minimize uncertainty and variability, which could be used by an agent to improve their control over the environment and increase extrinsic reward intake in the long run(40,41). Developments in reinforcement learning and robotics support the idea that both extrinsic and intrinsic rewards are important for maximizing an agent’s survival(42–44). Building intrinsic motivation into RL agents can promote the search for uncertain states and facilitate the acquisition of skills that generalize better across different environments, an essential feature for maximizing an agent’s ability to survive and reproduce over its lifetime, i.e. its evolutionary fitness(42).

The intrinsic reward of choice may be a specific instance of more general motivational constructs such as autonomy(13,14), personal causation(17), effectance(18), learned helplessness(45), perceived behavioral control(19) or self-efficacy(15), which are key for motivating behaviors that are not easily explained as satisfying basic drives such as hunger, thirst, sex, or pain avoidance(20). Common across these theoretical constructs is that control is intrinsically motivating only when the potential exists for agents to determine their own behavior, which when realized can give rise to a sense of agency and, in turn, strengthens the belief in the ability to exercise control over one’s life(46). Thus, individuals with an *internal* locus of control tend to believe that they, as opposed to external factors such as chance or other agents, control the events that affect their lives. Crucially, the notion of locus of control makes specific predictions about the relationship between preference for choice—choice being an opportunity to exercise control—and the environment: individuals should express a weaker preference for choice when the environment is adverse, stressful or unpredictable(47). This prediction is consistent with what is known about the influence of environmental adversity on control externalization: individuals exposed to greater environmental instabilities tend to believe that external and uncontrollable forces are the primary causes of events that affect their lives, as opposed to themselves(48). In other words, one would expect belief in one’s ability to control events, and thus preference for choice, to decline as the environment is perceived as increasingly unpredictable.

In our third experiment, we sought to test whether it was specifically a belief in personal control that motivated subjects, by altering the perceived controllability of the task environment. To do so, we introduced a novel change to the two-stage task where in a fraction of trials subjects experienced random swapping of the terminal states (fractals). Thus, subjects were subjected to trials where the terminal state was incoherent with their choice, and thus experienced alterations in their ability to predict the state of the environment following their action. Incoherence occurred with equal probability following free and forced actions in order to equate for any value associated with swapping itself. We found a significant reduction in the propensity to switch from forced to free choice following action-target incoherence, suggesting that altering the perceived controllability of the task causes choice to lose its attractiveness. This reduction in choice preference following incoherent trials is reminiscent of a form of locus externalization, and is consistent with the notion that choice preference is driven by a belief in one’s personal control. In this experiment, we focused on the value of personal control, and therefore held other decision variables such as outcome diversity(33,34), outcome entropy(35), and instrumental divergence (36,49). Further experiments are needed to understand how these variables interact with personal control in the acquisition of potential control over the environment.

Interestingly, when subjects selected the *free* option, the subsequent choice was sensitive to the past reward when the terminal state (the selected target) was coherent and the reward could therefore be attributed to the subject’s action. In contrast, subjects’ choices were insensitive to past reward when the terminal state was incoherent. Furthermore, the probability of sticking with the previous 2^nd^-stage choice following incoherent trials, whether rewarded or not, was not different from the probability of sticking with the previously *rewarded* 2^nd^-stage choice following coherent trials. Thus, subjects appeared to ignore information about action-state-reward contingencies that was externally derived, and instead appeared to double down by repeating their past choice as if they sought to maintain or regain personal control. This behavior is consistent with many observations suggesting that when individuals experience situations that threaten or reduce their personal control, they implement compensatory strategies to restore their perceived control to its baseline level(50,51).

Computationally, however, this compensatory strategy is at odds with a pure model-based strategy(37), where an agent could exploit information about action-state-reward contingencies whether it derived from their own choices (internal control) or from the environment (external control). Rather, it is consistent with work showing that choice-seeking could emerge when self-determined actions amplify subsequent positive reward prediction errors(5,52), and more generally with the notion that events are processed differently depending on individuals’ beliefs about their own control abilities. Thus, positive events are amplified only when they are believed to be within one’s personal control, whereas they are treated impartially when they are not(52), or when they come from an uncontrollable environment(53).

Together, our results suggest that choice seeking may represent one critical facet of intrinsic motivation and is associated with the desire of personal control. They also suggest that the need for personal control can compete with maximization of extrinsic reward provided by externally driven actions. Indeed, subjects favor positive outcomes associated to internally driven action even if reward rate is lower than for action performed under the instruction of an external agent. In general, the perception of being in personal control could then account for several aspects of our daily life such as enjoyment during game(54) or motivation to perform demanding task(55). Since our results shown inter-individual difference, it would be nonetheless important in the future to phenotype subject behaviors during choice-making to investigate how these individual traits can explain attitude difference when facing decision and their consequences, as exemplified by the variety of attribution and explanation styles of individuals in the general population(56,57).

## Materials and Methods

### Participants

Ninety-four healthy individuals (mean age = 30 ±SD 7.32 years, 64 females) responded to posted advertisements and were recruited to participate in this study. Relevant inclusion criteria for all participants were being fluent in French, not treated for neuropsychiatric disorders, having no color vision deficiency and being aged between 18 and 45 years old. Out of these 94 subjects, 58 participated to experiment 1 and 36 to experiments 2-3. We gave subjects 40 euros for participating. The sample size was chosen based on previous studies that used similar two-alternative decision making tasks(52,58,59).

### Ethics statement

The local ethics committee (Comité d’Evaluation Éthique de l’Inserm) approved the study (2019-CER2-MR-004). Participants gave written informed consent during inclusion in the study, which was carried out in accordance with the declaration of Helsinki (1964; revised 2013).

### General procedure

The paradigm was written in Matlab, using the Psychophysics Toolbox extensions(60,61). It was presented on a 24 inches screen (1920 × 1080 pixels, aspect ratio 16:9). Subjects seat ∼57 cm from the center of the monitor. Our behavioral task design was designed as a value-based decision paradigm. All participants received written and oral instructions. They were told that the goal of the task was to gain the maximum number of rewards (a large green euro). They were informed about the differences between the different trial types and that the extrinsic reward contingencies experienced during the training phases remained identical during the test phases. After instructions, participants received a pre-training session of a dozen trials (pre-train and pre-test phases) in order to familiarize them with the task design and the keys they would have to press.

In our experiments, subjects performed repeated trials with a two-stage structure. In the 1^st^-stage they made an initial decision about what could occur in the 2nd-stage. Selecting the *free* option led to a subsequent opportunity to choose and selecting the *forced* option led to an obligatory computer-selected action. In the 2^nd^-stage, we presented subjects with two fractal images, one of them being more rewarded following *free* selection in experiment 1 (except for P=0.5) and experiment 2. In experiments 1 and 2, the computer always selected the same fractal target following *forced* selection. Experiment 3 all fractal targets were equally rewarded and the computer randomly selected one of the two fractal targets following *forced* selection (50%). Following *forced* selection, the target to select with a key press was indicated by a grey V-shape above the target. Pressing the other key on this trial type did nothing and the computer waited for the correct key press to proceed further in the trial sequence. Either at the 1^st^- or 2^nd^-stage, after the subject’s selection of the target, a red V-shape appears immediately after above the target to indicate the one they had selected (in experiment 3 blocks this red V-shape remains 250ms on the screen and eventually jumped with the target, see below).

### Experimental conditions

In experiment 1, fifty-eight subjects performed trials where the maximal reward probabilities were matched following *free* and *forced* selection. We varied the overall expected value across different blocks of trials, each of them being associated to different programmed extrinsic reward probabilities (P). Forty-eight subjects performed a version with 3 blocks (experiment 1a) with different extrinsic reward probabilities ranging from 0.5 to 1 (block 1: P_*forced*_ = P_*free*_ = 0.5; block 2: P_*forced*_ = 0.75, P_*free*_|a1 = 0.75, P_*free*_|a2 = 0.25; block 3: P_*forced*_ = 1, P_*free*_| a1 = 1, P_*free*_|a2 = 0; where a1 and a2 represent the two possible key presses associated with the fractal targets). Ten additional subjects performed the same task with 4 different blocks (experiment 1b) associated to extrinsic reward probabilities also ranging from 0.5 to 1 (P = 0.5 or 0.67 or 0.83 or 1 from block 1 to 4 respectively.) We did not observe any substantial difference between these two subject groups, and pooled them for analyses.

Experiment 2 was similar to experiment 1 (six states) except programmed extrinsic reward associated with the *forced* option were higher than than the *free* option in two out of three blocks (P_*forced*_ = 0.75, 0.85 or 0.95). Reward probabilities following *free* selection did not change across the three blocks (P_*free*_|a1 = 0.75, P_*free*_|a2 = 0.25)

Experiment 3 consisted of a 7-state version of the two-stage task. Here, we manipulated the coherence between the subject selection of a 2^nd^-stage (fractal) target and the target ultimately displayed on the screen by the computer. Irrespectively of the target finally selected by the computer or the subjects, the extrinsic reward probability associated to all the 2^nd^-stage targets in *free* and *forced* trials was set at P=0.75. Importantly, adding the 7^th^ state in this last task version allowed the computer to swap the fractal 2^nd^-stage targets following both *free* and *forced* selection. Thus, subjects did not perceive the weak coherence as a feature specific to the *free* condition.

We associated unique fractal targets with each action in the 2^nd^-stage, and a new set was used for each block in all experiments. Colors of the 1^st^-stage targets were different between experiments. Positive or negative reward feedback, as well as the side of the 1^st^-stage and 2^nd^-stage target positions, were pseudo-randomly interleaved on the right or left of screen center. Feedback was represented by the presentation (reward) or not (non-reward) of a large green euro image.

In experiment 1, when P<1, participants performed a minimum of 48 trials per block in the training phases (*forced* and *free*) and the test phases. For P=1, participants performed a minimum 24 trials for training phases (*forced* and *free*) and 48 trials for test phase. The order of the blocks were randomly interleaved. In experiments 2 and 3 they performed a minimum of 40 trials for each block. Here, subjects started by performing experiment 3 followed by experiment 2. This was to ensure that the value of *free* trials was not devalued by experiment 2 (titration) when performing experiment 3. In experiment 3, subjects always started by the block with no target swaps (incoherence = 0), and in experiment 2 by the block with equal extrinsic reward probability (equivalent to the block P=0.75 of experiment 1). All the other blocks were randomly interleaved.

### Trial structure

During the training phase, for each trial, a first fixation point appeared in the center of the screen for 500ms, followed by the one of the first two targets of the different trial types for an additional 750ms, either (*forced* or *free*) to the left or right of the fixation point (∼11° from the center of the screen on the horizontal axis, 3° wide). Immediately after, the first target was turned off and two fractal targets appeared at the same eccentricity than the first target to the left and right of the fixation point. The subjects could then choose by themselves or had to match the target (depending on the trial type) using a key press (left or right arrow keys for left and right targets, respectively). After their selection, a red V-shape appeared for about 1000ms above the selected target (trace epoch). Note that in experiment 3, the V-shape was initially light red and turned on for 250ms above the actual fractal target selected by the subjects. It was then turn in dark red for 750ms. If the trial was incoherent, after 250ms, the red V-shape jumped and thus reappeared simultaneously with the other target on the other side of the screen also for 750ms. Finally, the fixation point was turned-off and the outcome was displayed during 750ms before the next trial. For the test phase, the timing was equivalent except for the decision epoch related to the first stage where participants could choose their favorite trial type (*free* and *forced* targets positioned randomly, left or right) after 500ms of fixation point presentation. When a selection was made, the first target remained for 500ms, associated to a red V-shape over the selected 1^st^-stage target – indicating their choice. The second stage started with a 500ms epoch where only the fixation point was presented on the screen, followed by the fractal target presentation. During the first and second action epochs, no time pressure was imposed on subjects to make their choice, but if they pressed one of the keys during the first 100ms after target presentation (‘early press’), a large red cross was displayed in the center of the screen for 500ms and the trial was repeated.

### Computational modelling

We fitted individual subject data with variants of temporal-difference reinforcement learning (TDRL) models. All models maintained a look-up table of state-action value estimates (*Q*(*s, a*)) for each unique target and each action across all conditions within a particular experiment. State-action values were updated at each stage (*t* ∈ {1,2}) within a trial according to the prediction error measuring the discrepancy between obtained and expected outcomes:

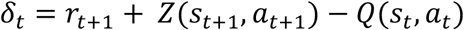

where *r*_*t*+1_ ∈ {0,1} indicates whether the subject received an extrinsic reward, and *Z*(*s*_*t*+1_, *a*_*t*+1_) represents the current estimate of the state-action value. The latter could take three possible forms:

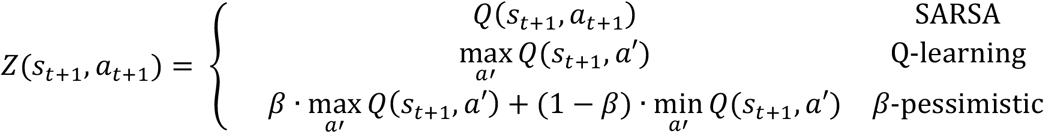

Although Q-learning and SARSA variants differ in whether they learn off- or on-policy, respectively, we treated both of these algorithms as optimistic. Q-learning is strictly optimistic by considering only the best future state-action value, whereas SARSA can be more or less optimistic depending on the sensitivity of the mapping from state-action value differences to behavioral policy. We compared Q-learning and SARSA variants with a third state-action value estimator that incorporates risk attitude through a weighted mixture of the best and worst future action values (Gaskett’s *β*-pessimistic model(32)). As *β* ⟶ 1 the pessimistic estimate of the current state-action value converges to Q-learning.

The prediction error was then used to update all state-action values according to:

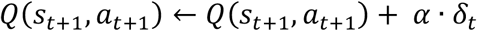

where *α* ∈ [0,1] represents the learning rate.

We tested whether a free choice bonus could explain choice preference by modifying the obtained reward as follows:

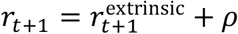

where *ρ* ∈ (−inf, +inf) is a scalar parameter added to any extrinsic reward following any action taken following selection of the free option.

Free actions at each stage were generated using a softmax policy as follows:

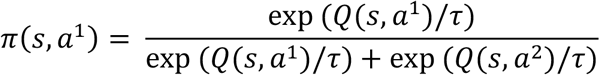

where increasing the temperature, *τ* ∈ [0, +inf), produces a softer probability distribution over actions. The forced option, on the other hand, always led to the same fixed action. We used a softmax behavioral policy for all TDRL variants, and in the context of our task, the Q-learning and SARSA algorithms were often similar in explaining subject data, so we treated them together in the main text (Supplementary Fig. 1).

We also tested the possibility that subjects exhibited tendencies to alternate or perseverate following free or forced actions. We implemented this using a stickiness parameter that modified the policy as follows:

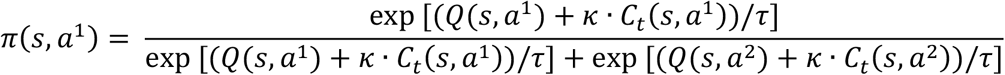

where the *κ* ∈ (−inf, +inf) parameter represents the subject’s tendance to perseverate, and *C*_*t*_(*s, a*) is a binary indicator for which fractal and action was chosen on the previous trial.

We independently combined the free parameters to produce a family of model fits for each subject. We allowed the learning rate (*α*) and softmax temperature (*τ*) to differ for each of the two stages in a trial. We therefore fitted a total of 48 models (3 estimates of current state-action value [SARSA, Q, *β*-pessimistic] × presence or absence of free choice bonus [*ρ*] × 2- vs 1-learning rate [*α*] × 2- vs 1-temperature [*τ*] × presence or absence of stickiness [*κ*]).

### Parameter estimation and model comparison

We fitted model parameters using maximum a posteriori (MAP) estimation using the following priors:

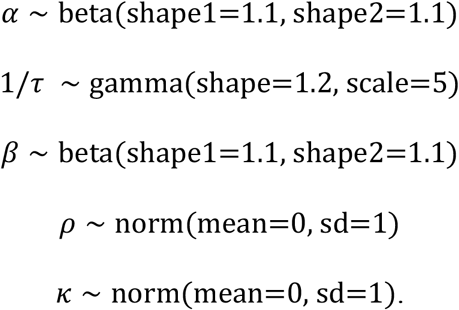

We based hyperparameters for *α* and 1/*τ* on Daw and colleagues (37). We used the same priors and hyperparameters for all models containing a particular parameter. We used limited-memory quasi-Newton algorithm (L-BFGS-B) to numerically compute MAP estimates, with *α* and *β* bounded between 0 and 1 and 1/*τ* bounded below at 0. For each model, we selected the best MAP estimate from 10 random parameter initializations.

For each model for each subject, we fitted a single set of parameters to both training and test data across conditions. We initialized state-action values to zero at the beginning of the training phase for each condition. Data from the training phase consisted of 2^nd^-stage actions and rewards, but we also presented subjects with the 1^st^-stage cues corresponding to the condition being trained (forced or free). Therefore, we fitted the TDRL models assuming that the state-action values associated with the 1^st^-stage fractals also underwent learning during the training phase, and that these backups continued into the test phase, where subjects actually made 1^st^-stage decisions. That is, we initialized the state-action values during the test phase with the final state-action values during the training phase.

We used Schwarz weights to compare models, which provides a measure of the strength of evidence in favor of one model over others and can be interpreted as the probability that a model is best in the Bayesian Information Criterion (BIC) sense(62). We calculated weights for each model as:

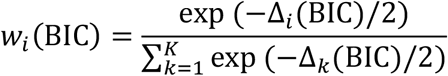

so that ∑*w*_*i*_ (BIC) = 1. We selected the model with the maximal Schwarz weight for each subject.

In order to verify that we could discriminate different state-action value estimates and how accurately we could estimate parameters, we performed model and parameter recovery analyses on simulated datasets (Supplementary Fig. 1).

### Statistical analyses

We used generalized linear mixed models (GLMM) to examine differences in choice behavior. When the model did not include trial-specific information (e.g., reward on the previous trial), we aggregated data to the block level. Otherwise, we used choice data at the trial level. We included random effects by subject for all models (random intercepts and random slopes for the variable manipulated in each experiment; maximal expected value, relative expected value, or incoherence for experiments 1, 2, and 3, respectively). We performed GLMM significance testing using likelihood-ratio tests, and we corrected for multiple comparisons in post-hoc tests using Tukey’s method. We used generalized additive mixed models (GAMM) to examine choice behavior as a function of trial within a block. We obtained smooth estimates of choice behavior using penalized regression splines, with penalization that allowed smooths to be reduced to zero effect(63). We included separate smooths by block. We performed GAMM significance testing using approximate Wald-like tests(64).

## Acknowledgements

J.M. was supported by the Agence Nationale de la Recherche (ANR) grant ANR-19-CE37-0014-01 (ANR PRC) and by the European Commission (H2020-MSCA-IF-2018-#845176). D.B. was supported by a FRM fellowship (FDM201906008526). V.C. was supported by the ANR grants ANR-17-EURE-0017 (Frontiers in Cognition), ANR-10-IDEX-0001-02 PSL (program ‘Investissements d’Avenir’), ANR-16-CE37-0012-01 (ANR JCJ) and ANR-19-CE37-0014-01 (ANR PRC). B.L. was supported by the ANR grant ANR-19-CE37-0014-01. The authors of this article are grateful to Karim Ndiaye, operational manager of the PRISME platform at the ICM for his valuable help during participant testing.

## Author Contributions

J.M., V.C. and B.L. designed the study; J.M., M.R.A., D.B. and A.K. performed the experiments and preliminary analyses V.C.; J.M., and B.L. designed and performed final analyses; J.M., V.C. and B.L. wrote the manuscript.

## Data availability statement

All data and related metadata underlying the findings reported will be deposited in Zenodo (DOI: 10.5281/zenodo.7057043) at the time of publication.

## Code reporting

Code written in support of this publication will be made publicly available in Zenodo (DOI: 10.5281/zenodo.7057080) at the time of publication.

