## Supplemental information for "Choice seeking is motivated by the intrinsic need for personal control"

**This PDF file includes:**

Supporting text
Figures S1 to S3
SI References

**Supporting Information Text**

In order to verify that we could discriminate different TDRL models and accurately estimate parameters, we performed model and parameter recovery analyses on simulated datasets. We considered six key models, the three different value estimates (SARSA, Q-learning, $\beta$-pessimistic) with and without a free choice bonus ($\rho$). For each of these models, we simulated the performance of 5000 virtual subjects using the same set of conditions that real subjects experienced in experiment 1. We explored a wide parameter space, drawing generative parameters from the following distributions:

$$\alpha \sim \text{beta(shape1=1.75, shape2=2.75)}$$
$$1/\tau \sim \text{gamma(shape=2.5, scale=1)}$$
$$\beta \sim \text{beta(shape1=1, shape2=2.25)}$$
$$\rho \sim \text{gamma(shape=2.5, scale=0.25)}$$

We fitted simulated data and performed model selection using the same procedures as for real data. From this data, we calculated confusion and inversion matrices(1) (Fig. S1A), which give the probability that the data simulated by one model is best fit by another and the probability that one model is more likely generated by another, respectively. We found that model recovery was good, although Q-learning and SARSA algorithms were confused in the parameter ranges used. This is due to the fact that Q-learning is strictly optimistic by considering only the best future state-action value, whereas SARSA can be more or less optimistic depending on the sensitivity of the mapping from state-action value differences to behavioral policy. Thus, our use of a softmax policy means that SARSA approaches Q-learning as $1/\tau$ increases. Therefore, we treated Q-learning and SARSA algorithms together as "optimistic" in the main text. Pooling these two models together indicates that model recovery is good (Fig. S1B).

We also examined parameter recovery using correlations between estimated and simulated parameters, both with the same parameter as well as across parameters (Fig. S1C-D). We found that parameter recovery was good (diagonal correlations), and parameter confounding was quite weak (off-diagonal correlations).

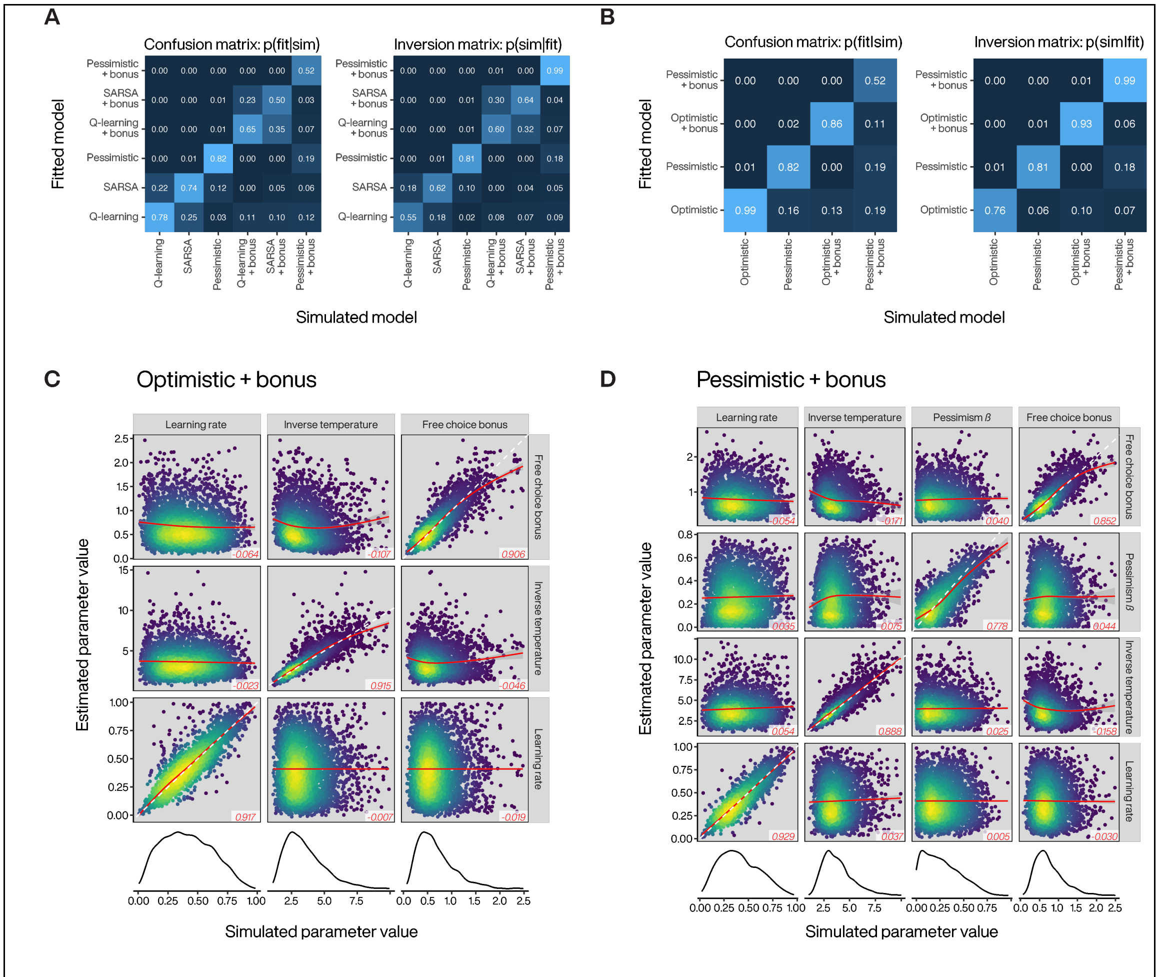


**Figure S1.** Model and parameter recovery simulations. **A.** Confusion and inversion matrices for six models varying value update target (Q-learning, SARSA, or $\beta$-pessimistic) and free choice bonus (presence or absence). **B.** Confusion and inversion matrices formed by treating Q-learning and SARSA together as models with "optimistic" value update targets. **C.** Point density plots of simulated and estimated parameter values for models with an optimistic target (both Q-learning and SARSA). The dashed white line in the diagonal panels has unity slope, and the red lines represent an additive model smooth. The red numbers listed in the lower right of each panel are the correlation coefficients for each simulated and estimated parameter. The bottom row contains density estimates for the simulated parameter values. **D.** As C. for $\beta$-pessimistic target with a free choice bonus.

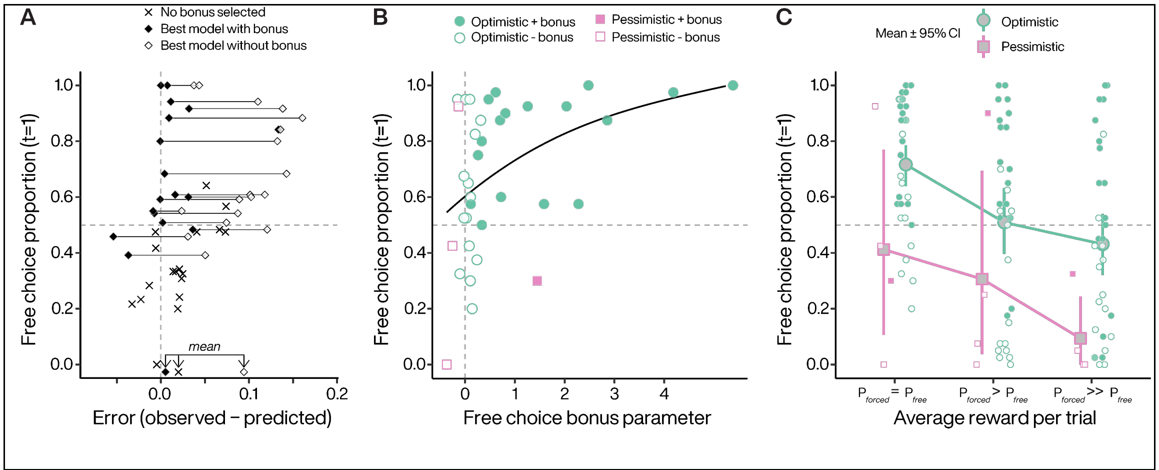


**Figure S2.** Reinforcement learning models capture individual choice behavior in experiment 2. **A.** Obtained free choice proportion as a function of model error, averaged over all conditions. For subjects where the selected model did not include a free choice bonus, only one symbol (X) is plotted. For subjects where the selected model included a free choice bonus, two symbols are plotted. Filled symbol represents the fit error with the selected model, and the open symbol represents the next best model that did not include a free choice bonus. Lines connect individual subjects. **B.** Bonus coefficients increase as a function of subjects' preference for *free* options irrespectively of the target policy they used when performing the task. Choice preference from equal probability blocks (P=0.75). Filled circles indicate that the best model included a free choice bonus parameter. Line illustrates a generalized additive model smooth. **C.** Both optimistic and pessimistic subjects decrease their *free* option preference as the *forced* option value increases. Symbol legend from B applies to the small points representing individual means in C. Error bars for 95% CI.

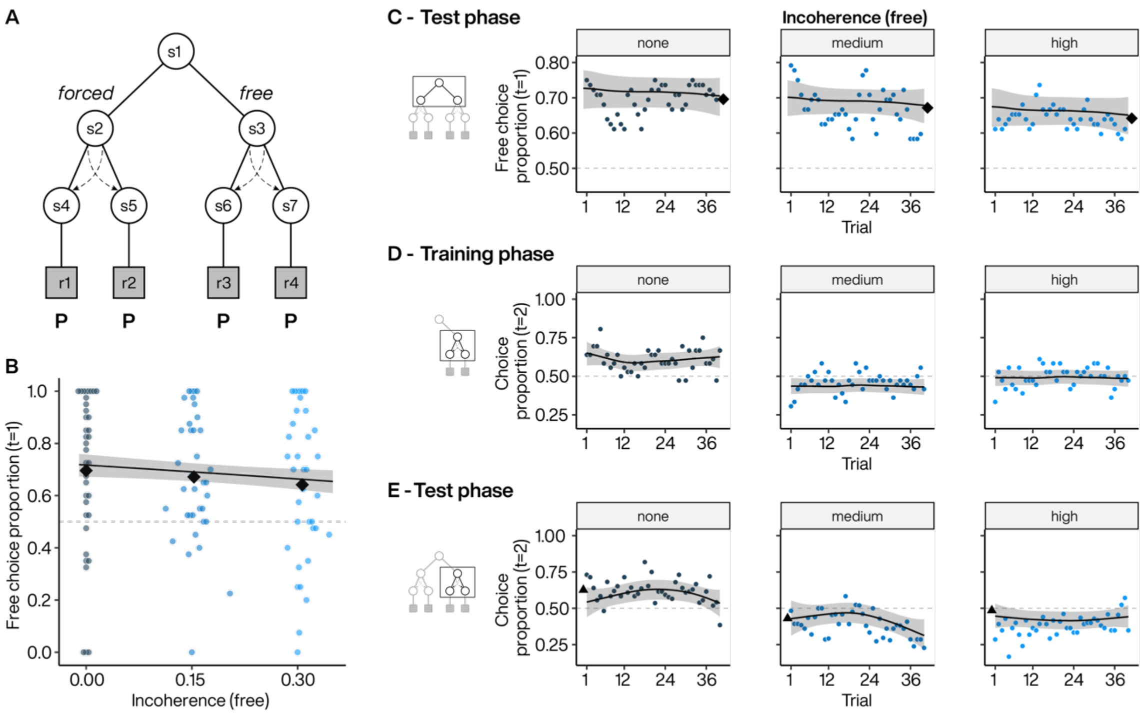


**Figure S3.** Choice proportion across different action-outcome incoherence level. A. Experiment 3 task design using seven states task design where we manipulated the incoherence in both free and forced options. B. Subject preference for free options during 1st-stage. Colored points indicate individual subject mean choice preference per block, plotted against the incoherence level. Black diamonds indicate the average of subject means per block. Line indicates the estimated choice preference from a GLMM, with 95% CI. C. Dynamics of free option preferences across test phase blocks for incoherence set at 0 (i.e., none, left), 0.15 (i.e. medium, middle) and 0.30 (i.e. high, right). Each point represents the average free option preference as a function of trial within a block. Diamonds: as in B. Lines indicate the estimated choice preference from a GAMM, with 95% CI. D to E. Dynamics of the selection of the two 2nd-stage targets (equally rewarded) in free options across the blocks for incoherence levels set at 0 (left), 0.15 (middle) and 0.30 (right) during the training (D) and test (E) phases. Triangles represents the final average selection at the end of the training phases. Lines: as in C.
